# Pre-copulatory reproductive behaviours are preserved in *Drosophila melanogaster* infected with bacteria

**DOI:** 10.1101/2021.09.15.460306

**Authors:** Saloni Rose, Esteban J. Beckwith, Charlotte Burmester, Robin C. May, Marc S. Dionne, Carolina Rezaval

## Abstract

Reproduction and immunity are crucial traits that determine an animal’s fitness. Terminal investment hypothesis predicts that reproductive investment should increase in the face of a mortality risk caused by infection. However, due to competitive allocation of energetic resources, individuals fighting infections are expected to decrease reproductive efforts. While there is evidence for both hypotheses, the factors that determine the choice between these strategies are poorly understood. Here, we assess the impact of bacterial infection on pre-copulatory behaviours in the fruit fly *Drosophila melanogaster*. We found that male flies infected with six different bacteria, including pathogenic and non-pathogenic strains, show no significant differences in courtship intensity and mating success. Similarly, bacterial infections did not affect sexual receptivity in female flies. Our data suggest that pre-copulatory reproductive behaviours remain preserved in infected animals, despite the huge metabolic cost of infection.

## Introduction

Reproduction and immunity are intricately linked traits central to an animal’s fitness [1]. Lifehistory theory suggests that there is a trade-off between these two processes as the energy reserves are limited [2]. In the face of infection, animals need to mount an immune response that is energetically expensive [3,4]. As a result, infected animals might invest less in reproduction. Previous studies in insects and birds have shown that egg production can decrease upon infection [1,5]. For example, *Anopheles* females infected with the malarial parasite *Plasmodium* or injected with lipopolysaccharide (LPS), a bacterial cell wall component, produce a lower number of eggs [6,7]. In males, immune system activation has been shown to reduce sperm production and viability in crickets, birds and fish [8]. The decrease in reproductive investment is not limited to postcopulatory traits. Some studies report reduced expenditure in pre-copulatory reproductive behaviours in response to infection. For example, Charge et al. showed that immune challenged male houbara bustards (*Chlamydotis undulata*) display reduced courtship behaviours [9]. As the animal recovers from the infection, the courtship levels return to normal. Similarly, fishes infected with ectoparasites such as *G. turnbulli* or endoparasites such as microsporidians have been shown to invest less time in courtship [10–12]. These studies argue that individuals exposed to harmful infections prioritise defence over reproduction investment, thus increasing the likelihood of survival [13–16].

In an alternative, and opposite, scenario, the terminal investment hypothesis predicts that, upon perceiving a self-integrity or mortality risk, an individual will increase reproductive effort as its chance for future reproduction decreases. Through this strategy, individuals would shift their investment away from defence and repair traits and maximise their lifetime reproductive success [17]. Studies in diverse species have provided evidence for terminal investment. For example, Japanese tree frogs increase their calling effort to attract a mate in response to chytrid fungus infection [18]. Similarly, desert locusts (*Schistocerca gregaria*)infected with *Metarhizium acridum*, an entomopathogenic fungus, display increased mounting behaviour [19]. Male crickets (*Allonemobius socius*) injected with LPS employ different strategies based on their age since old, but not young, males accelerate their reproductive efforts upon infection [20]. Therefore, studies across taxa have found support for both reproduction-immunity trade-off and terminal investment hypothesis. This prompts the interesting question: what factors determine the selection of one investment strategy over the other? Strategies for balancing immunity and reproductive efforts may vary depending on the host’s internal conditions (e.g., age and genetic background)[20,21] and extrinsic conditions (e.g., nature of the infection, pathogen’s virulence, and its relationship with the host) [22]. Investigating how the infection type, timing, and virulence modulates reproductive behaviours is essential for understanding variation in reproductive effort across species.

The use of genetically tractable experimental systems opens the possibility of uncovering the genetic determinants of this process. *Drosophila melanogaster* is a powerful model organism for behavioural ecology and neuroscience [23–25] and is very well suited to study the interaction between reproduction and immunity. Fruit flies display an innate courtship ritual that allows them to evaluate the suitability of a potential mate while increasing the chances of successful copulation [26]. While males display a series of stereotyped courtship behaviours, such as singing a species-specific song by vibrating a wing, female flies show acceptance or rejection behaviours in response to the male’s courtship display [26]. During this encounter, flies exchange olfactory, gustatory, and visual cues that signal sex, species, and mating status [26]. The circuitry underlying courtship is defined by the sex determination genes *doublesex* and *fruitless* and has been extensively studied [26]. Studies report that male fruit flies have a high level of sexual drive and courtship commitment, even in the presence of suboptimal mating targets [27,28]. However, sustained sexual activity for extended periods of time can be metabolically costly for the animal [29] and increase the chances of predatory risk.

With a well-characterised immune system, *Drosophila* can be easily infected with pathogens either naturally or in experimental situations and serves as an excellent platform to study the effects of infectious diseases. The fly immune system relies on multiple defences, including physical barriers, local responses such as the production of reactive oxygen species (ROS), a cellular response, and a systemic response characterised by secretion of antimicrobial peptides (AMPs) into the hemolymph to fight microbial pathogens [30,31]. While a hallmark of the immune activation is the synthesis of AMPs, it is characterized by a marked transcriptomic switch with a profound metabolic impact [32–35]. The AMP production and the metabolic switch takes place primarily in the fat body and is under the control of two different cascades: Toll and Imd pathways, which activate different NF-κB-like factors and induce the innate immune response. The Toll pathway responds to antigens from fungi and Gram-positive bacteria while the Imd pathway detects Gram-negative bacteria [30,31]. Notably, many aspects of innate immunity are conserved between flies and mammals, such as NF-kB family transcription factors and signal transduction pathways [30,31], and the organization of the immune system is highly analogous [36]. A few studies have investigated the impact of pathogen infection on female sexual traits, like oviposition [37] and some aspects of male courtship [38]. However, how infection modulates reproductive behaviours in *Drosophila* remains poorly understood.

Here, we carried out detailed analyses of the effect of infections with diverse bacteria on fly male courtship behaviours and female sexual receptivity. We systematically tested the behavioural impact of infections with pathogenic (*S. marcescens, S. aureus* and *L. monocytogenes*) and non-pathogenic species (*Erwinia carotovora carotovora 15* (*ECC15*), *E. coli* and *M. luteus*), which are phylogenetically diverse, have different virulence and hostpathogen relationships, and impose differing fitness costs on flies [39–41]. Our findings indicate that, regardless of the species’ virulence and pathogen-host relationship, bacterial infection does not significantly affect male courtship behaviour and mating success in *Drosophila melanogaster*. Similarly, female sexual receptivity remained unaffected upon infection. Moreover, genetic activation of Toll and Imd immune pathways in males and female flies had no impact on courtship displays. Our study demonstrates that pre-copulatory reproductive behaviours remain preserved in infected flies despite the significant metabolic cost of infection.

## Results

### Non-pathogenic infections do not affect courtship behaviour of wild-type males

To reliably produce infection phenotypes and assess the consequences in behaviour, we used a nano-injector to deliver precise volumes of bacterial solution into the abdomen of flies [42]. As a starting point, we chose three different non-pathogenic bacteria: *ECC15, E. coli* and *M. luteus*, which activate the fly’s innate immune system without affecting lifespan [43–45]. Gram-negative bacteria *ECC15* and *E*. coli induce the Imd pathway, while the gram-positive bacterium *M. luteus* activates the Toll pathway. We confirmed the reproducibility of our infection protocols by assessing the fly’s survival rate in response to pathogenic infections (**Fig S1**). As expected, CS males infected with either ECC15, *E. coli* or *M. luteus* showed a survival rate similar to that of uninfected and sham-infected flies (injected with PBS solution) (**Fig S1A-C**). Given that flies rapidly clear non-pathogenic bacteria from their bodies [40,45,46], we carried out our behavioural experiments 5-6 hours post-infection, when the bacterial load is still detectable.

To assess the effects of non-pathogenic bacteria on male courtship behaviour, we performed single mating assays, where an infected male was paired with a healthy virgin female (**Fig 1A)**. As expected, uninfected and sham-infected males spent most of the time courting a female (Median Courtship Index (CI): Uninfected= 95.07, PBS= 90.61; **Fig 1B**). Interestingly, males infected with *ECC15* displayed comparable courtship levels to that of controls (*ECC15* CI= 77.16; **Fig 1B**). Similarly, *E. coli* and *M. luteus* infection had minimal effect on male courtship behaviour (Uninfected CI= 99.52, CI PBS= 83.33 and CI *E. coli* = 84.73; **Fig 1C**; CI Uninfected= 90.7, CI PBS= 84 and CI *M. luteus* = 80.45; **Fig 1D**). Most control males successfully copulated within one hour and we found that infected males show a similar rate of copulation (Mating success: Uninfected= 100%, PBS= 100% and *ECC15* = 95.24%, **Fig 1E**; Uninfected= 100%, PBS= 100% and *E. coli* = 100%, **Fig 1F**; Uninfected= 100%, PBS= 95.24% and *M. luteus* = 100%, **Fig 1G**). Next, we asked if we could generalise these findings to a second *Drosophila* strain. To this end, we selected wildtype Dahomey, an outbred strain isolated from Benin [47]. We found that, like CS, Dahomey males did not alter their courtship behaviour in response to infection (**Fig S2**). These results suggest that male courtship behaviours remain unaffected in response to non-pathogenic infections.

**Figure 1:**
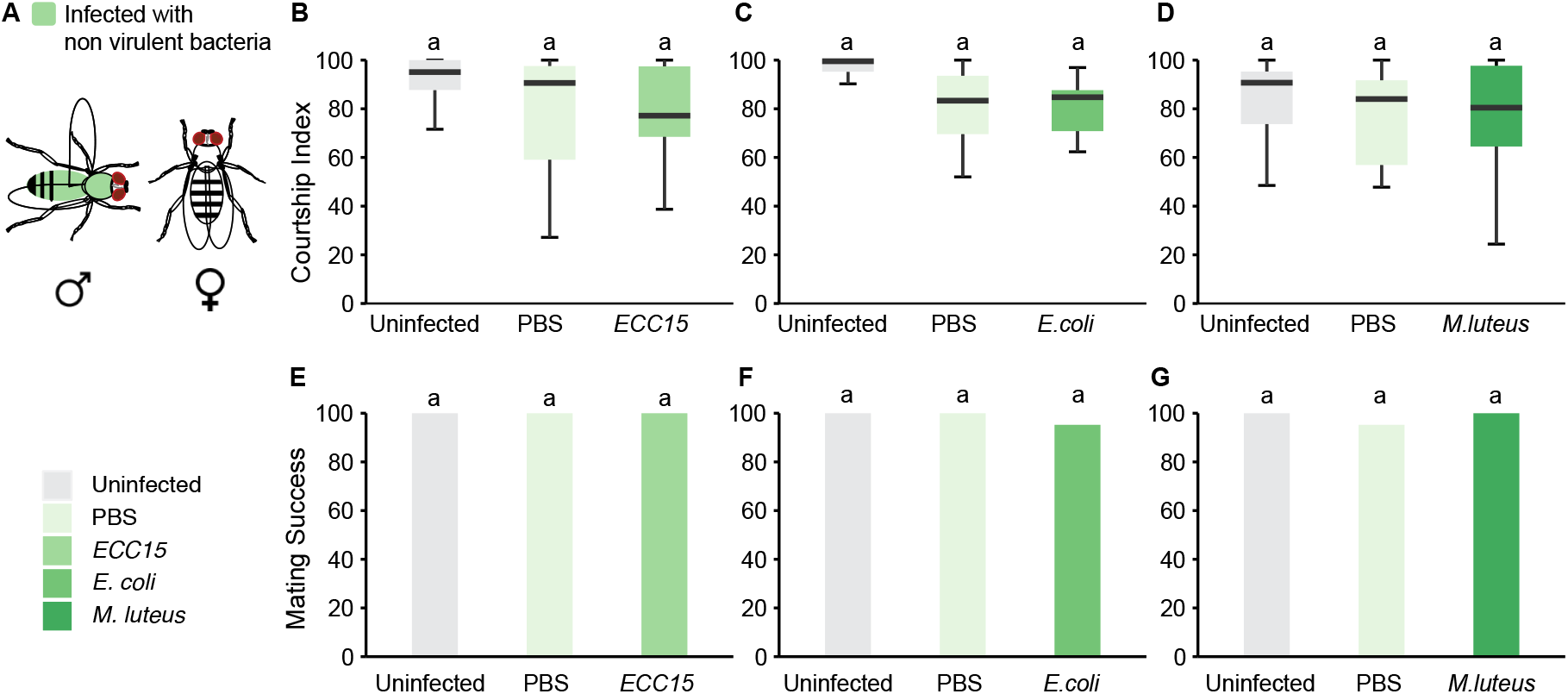
Effect of non-pathogenic bacterial infections on male courtship behaviour. (A) Male CS flies were injected with three different pathogens and tested in a single pair courtship assay with an uninfected virgin female. (B) Courtship index and (E) mating success of males infected with *ECC15* and their respective controls (n=20-21). (C) Courtship index and (F) mating success of males infected with *E. coli* and their respective controls (n=19-23). (D) Courtship index and (G) mating success of males infected with *M. luteus* and their respective controls (n=21-22). Dunn Test in BD and Fisher Test in E-G. No significant differences were observed between the treatments.

### Pathogenic infections do not affect courtship behaviour of wild-type males

Our findings show that infection with non-pathogenic strains do not reduce male courtship in males. We reasoned that more virulent and pathogenic bacteria that can reproduce within the flies may have a greater impact on male reproductive behaviours. To test this hypothesis, we chose three different pathogenic bacterial species that have been shown to negatively affect *Drosophila* physiology and induce lethality: *S. marcescens, S. aureus* and *L. monocytogenes*. Infecting male flies with the natural insect pathogen *S. marcescens* caused 100% mortality within 9 hours upon injection (**Fig S1D**). In contrast, uninfected flies and sham-infected flies survived through the observation time. Non-natural pathogens for flies, such as *S. aureus* or *L. monocytogenes*, induced lethality within 24 hours (**Fig S1E**) and ~7 days (**Fig S1F**) respectively, as previously reported [39,40,48]. On the basis of the survival data, we performed behavioural experiments with *S. marcescens, S. aureus* and *L. monocytogenes* at 5-6h, 8-9h, 24-25h post infection, respectively, a time period at which infection is advanced but not too detrimental for the flies (**Fig 2**).

**Figure 2.**
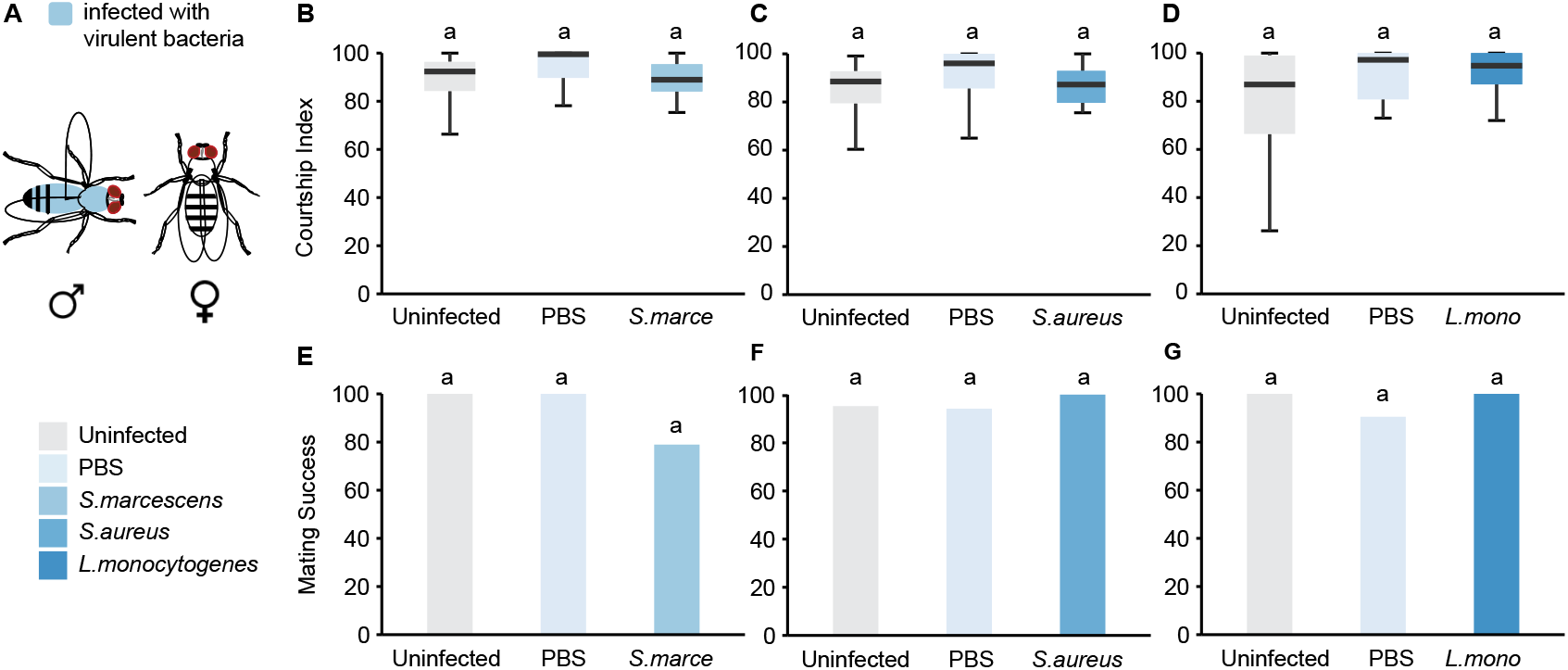
Effect of pathogenic bacterial infections on male courtship behaviour. (A) Male CS flies were injected with three different pathogens and tested in a single pair courtship assay with an uninfected virgin female. (B) Courtship index and (E) mating success of males infected with *S. marcescens* and their respective controls (n=18-19). (C) Courtship index and (F) mating success of males infected with *S. aureus* and their respective controls (n=17-21). (D) Courtship index and (G) mating success of males infected with *L. monocytogenes* and their respective controls (n=19-20). Dunn Test in B-D and Fisher Test in E-G. No significant differences were observed between the treatments.

Surprisingly, none of these lethal infections altered male courtship behaviour. There were no observable changes in the courtship index of infected males with either *S. marcescens* (Median CI: Uninfected= 92.35, PBS= 99.53 and *S. marcescens* = 88.94, **Fig 2B**), *S. aureus* (Median CI: Uninfected= 88.5, PBS= 96.04 and *S. aureus* = 87.25, **Fig 2C**) or *L. monocytogenes* (Median CI: Uninfected= 86.95, PBS= 97.19 and *L. monocytogenes* = 94.76, **Fig 2D**). Moreover, lethal infections did not compromise male mating success (Mating success: Uninfected= 100%, PBS= 100% and *S. marcescens* = 80%, **Fig 2E**; Uninfected= 95.24%, PBS= 94.12% and *S. aureus* = 100%, **Fig 2F**; Uninfected= 100%, PBS= 90.48% and *L. monocytogenes* = 100%, **Fig 2G**). Similarly, the courtship behaviours of Dahomey male flies infected with any of these pathogenic bacterial strains remained unaffected (**Fig S3**).

Altogether, our observations indicate that pathogenic and non-pathogenic infection does not affect or increase male courtship behaviours. We reasoned that the high level of sex drive and courtship index observed in control uninfected and sham-infected flies **(Fig 1 and Fig 2**) might mask any potential effects from bacterial infection. To decrease basal courtship levels, we placed CS males with mated females, which are reluctant to copulate and are therefore unattractive courtship targets [49]. As expected, uninfected CS males showed decreased courtship index towards mated females (~20%). However, neither pathogenic (*S. marcescens*) or non-pathogenic (*ECC15* and *M. luteus*) infections altered courtship behaviours (**Table S1**). Therefore, these findings reinforce our previous results showing that CS and Dahomey males infected with either pathogenic or non-pathogenic strains maintain their courtship efforts.

### Non-pathogenic infections do not affect sexual receptivity of wild-type females

Previous reports suggest that the response to infections is sexually dimorphic in many animals, including flies [50,51]. This dimorphism is seen in survival, pathology, bacterial loads and activity [50]. In addition, the costs of reproduction are different between sexes. Males invest considerable energy in courting a potential mate while females do not play an active role during courtship. Given that the costs of immunity and reproduction are different between males and females, we asked if females respond differently to infections. To test this, we injected CS and Dahomey virgin females with *ECC15, E. coli* and *M. luteus* and measured copulation latency and mating success, both of which are proxies for female receptivity (**Fig 3A**). Uninfected CS virgin females readily accept males and copulate within 2-5 minutes. We found that infected females and sham-infected females exhibited a higher latency to copulation than uninfected controls. However, there were no differences between sham-infected and infected females, indicating that bacterial infection itself does not influence copulation latency (Median Copulation Latency: Uninfected = 131s, PBS= 384.5s, *ECC15* = 402s, **Fig 3B**; Uninfected = 37.5 s, PBS= 500s, *E. coli* = 456s, **Fig 3C;** Uninfected = 248s, PBS= 567s; *M. luteus* = 574s, **Fig 3D**). Similarly, female mating success remained unchanged upon infection (Mating success: Uninfected= 95.83%, PBS= 100% and *ECC15=*100%, **Fig 3E**; Uninfected= 100%, PBS= 85% and *E. coli* = 89.47%, **Fig 3F**; Uninfected= 100%, PBS= 100% and *M. luteus* = 100%, **Fig 3G**). Dahomey virgin females infected with non-pathogenic strains showed a similar trend (**Fig S4**). We therefore conclude that non-pathogenic infections do not affect female receptivity in CS and Dahomey *Drosophila* females.

**Figure 3.**
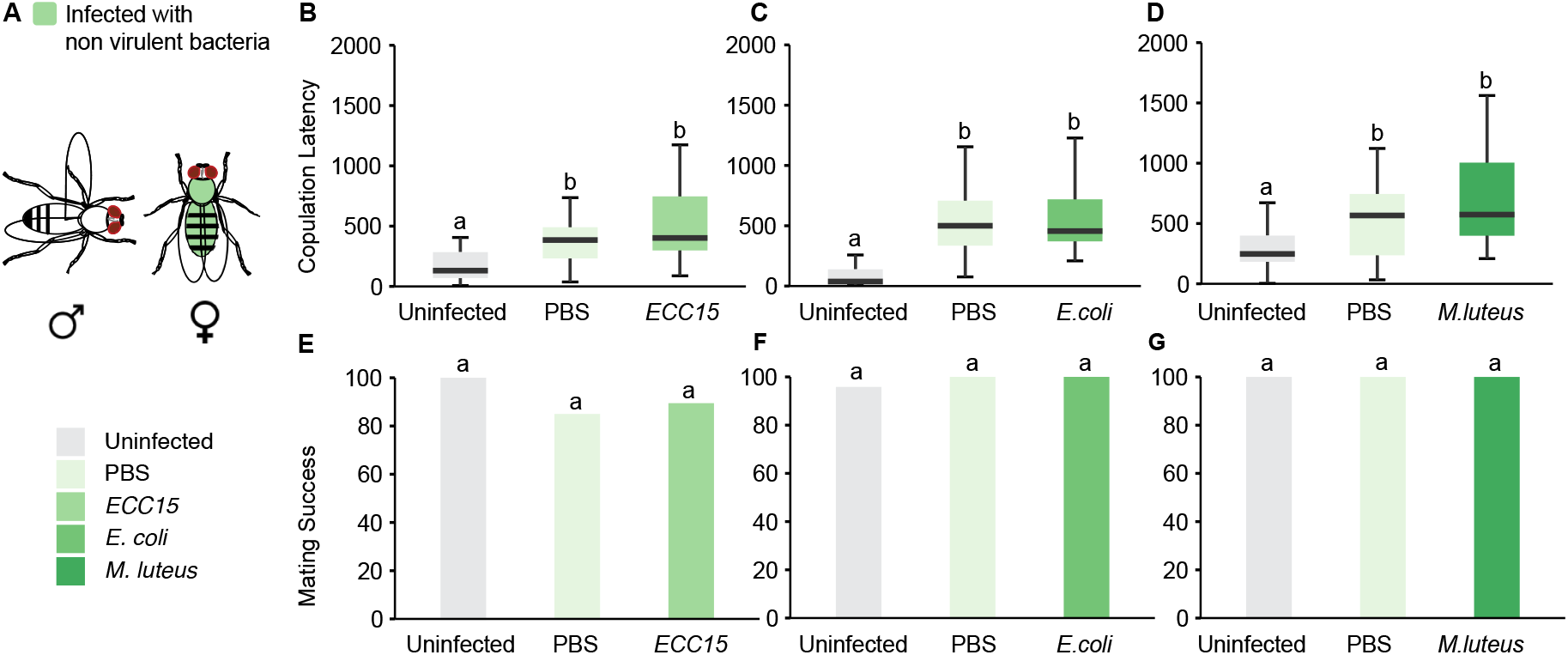
Effect of non-virulent bacterial infections on female receptivity. (A) Virgin female Canton S flies were injected with three different pathogens and tested in a single pair courtship assay with an uninfected male. (B) Copulation latency and (E) mating success of females infected with *ECC15* and their respective controls (n=20-24). (C) Copulation latency and (F) mating success of females infected with *E. coli* and their respective controls (n=20-21). (D) Copulation latency and (G) mating success of males infected with *M. luteus* and their respective controls (n=19-20). Dunn Test in B-D and Fisher Test in E-G.

### Pathogenic infections do not affect sexual receptivity of wild-type females

Next, we tested the effects of pathogenic strains *S. marcescens, S. aureus* and *L. monocytogenes* in female behaviour. We found that these infections dramatically reduced female survival (Fig S1). To test if female flies change their reproductive efforts in response to pathogenic infection, we injected CS virgin females with these pathogenic strains and presented them with a virgin CS male, which is sexually mature and has a high sex drive (**Fig 4A)**. Similar to males, infection status did not affect latency to copulation in females (Median Copulation Latency: Uninfected = 167s, PBS= 250.5s, *S. marcescens* = 312s, **Fig 4B**; Uninfected = 198.5s, PBS= 390s, *S. aureus* = 306s, **Fig 4C;** Uninfected = 146s, PBS= 98.5s; *L. monocytogenes* = 186.5s, **Fig 4D**). In addition, female mating success remained unaffected upon pathogenic infection (Mating success: Uninfected= 100%, PBS= 100% and *S. marcescens* = 81.25%, **Fig 4E**; Mating success: Uninfected= 94.63%, PBS= 95% and *S. aureus* = 100%, **Fig 4F**; Uninfected= 95.24%, PBS= 100% and *L. monocytogenes* = 95.24%, **Fig 4G**). We confirmed these findings using WT Dahomey females, where we observed no significant changes in copulation latency or mating success (**Fig S5**). Hence, pathogenic bacterial infections do not modulate sexual receptivity in *Drosophila* females.

**Figure 4.**
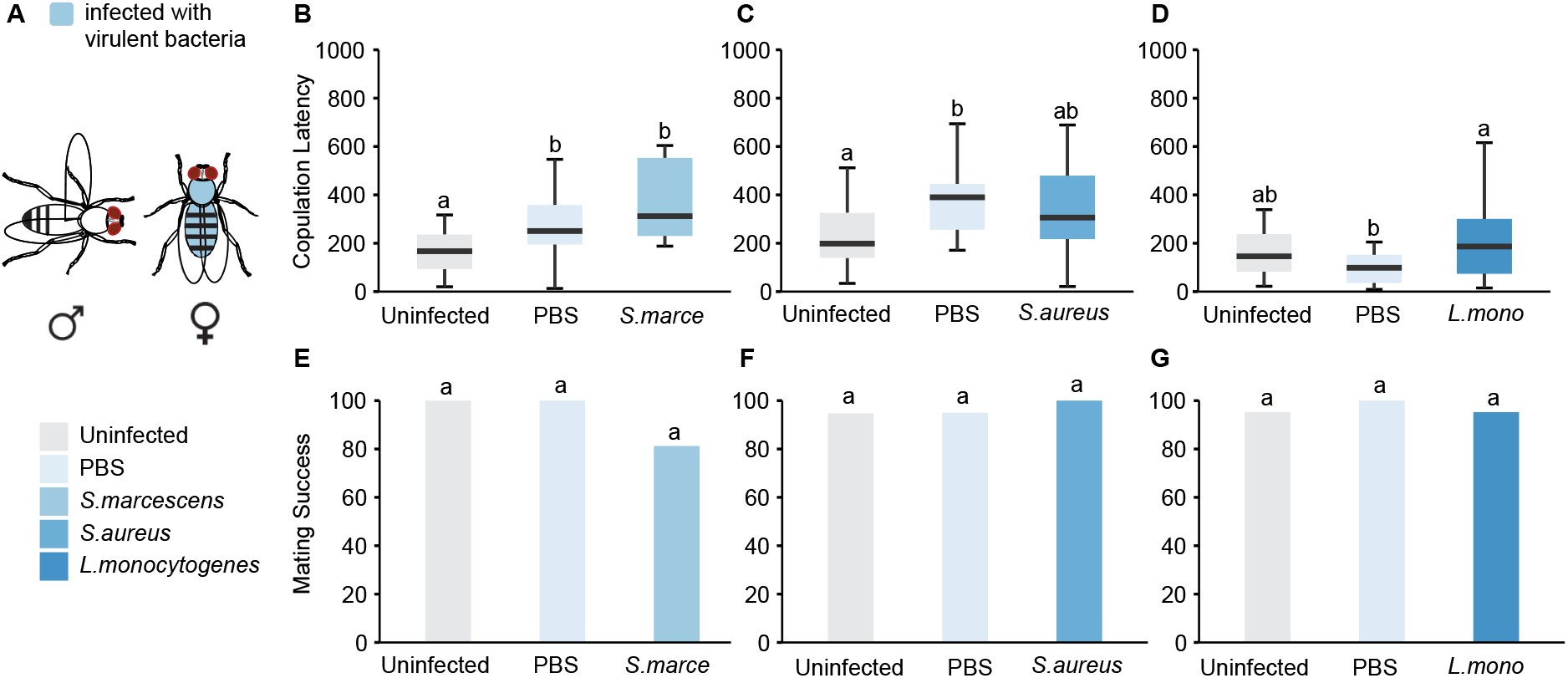
Effect of virulent bacterial infections on female receptivity. (A) Virgin female Canton S flies were injected with three different pathogens and tested in a single pair courtship assay with an uninfected male. (B) Copulation latency and (E) mating success of females infected with *S. marcescens* and their respective controls (n=16-19). (C) Copulation latency and (F) mating success of females infected with *S. aureus* and their respective controls (n=19-20). (D) Copulation latency and (G) mating success of males infected with *L. monocytogenes* and their respective controls (n=19-20). Dunn Test in B-D and Fisher Test in E-G.

### Effect of social context on pre-copulatory behaviours under infection conditions

Altogether, our data indicate that infections with different virulent and non-virulent pathogens do not modulate male courtship behaviours or mating success in *Drosophila*. We next wondered whether infected male flies would behave differently in the presence of a healthy male competitor (**Fig 5A**). To test this, we paired a CS virgin female with a healthy male and an infected male with *S. aureus*. Surprisingly, we found that infected males with this virulent pathogen courted to the same extent as the uninfected male (**Fig 5 B**). In addition, healthy females did not preferentially mate with healthy males over infected males (**Fig 5C)**. We similarly tested whether healthy male flies would behave differently towards healthy or infected females (**Fig 5D**). When given a simultaneous choice between *S. aureus* infected females and sham-infected females, these males spent a similar amount of time courting each female in a competitive assay (**Fig 5E**). Moreover, they mated equally with healthy *S. aureus*-infected females **(Fig 5F**). Extending these studies to a second virulent pathogen, *S. marcescens*, showed an identical pattern (**Fig 5G-L**). These results suggest that a more complex social context does not change the reproductive performance of sick flies.

**Figure 5.**
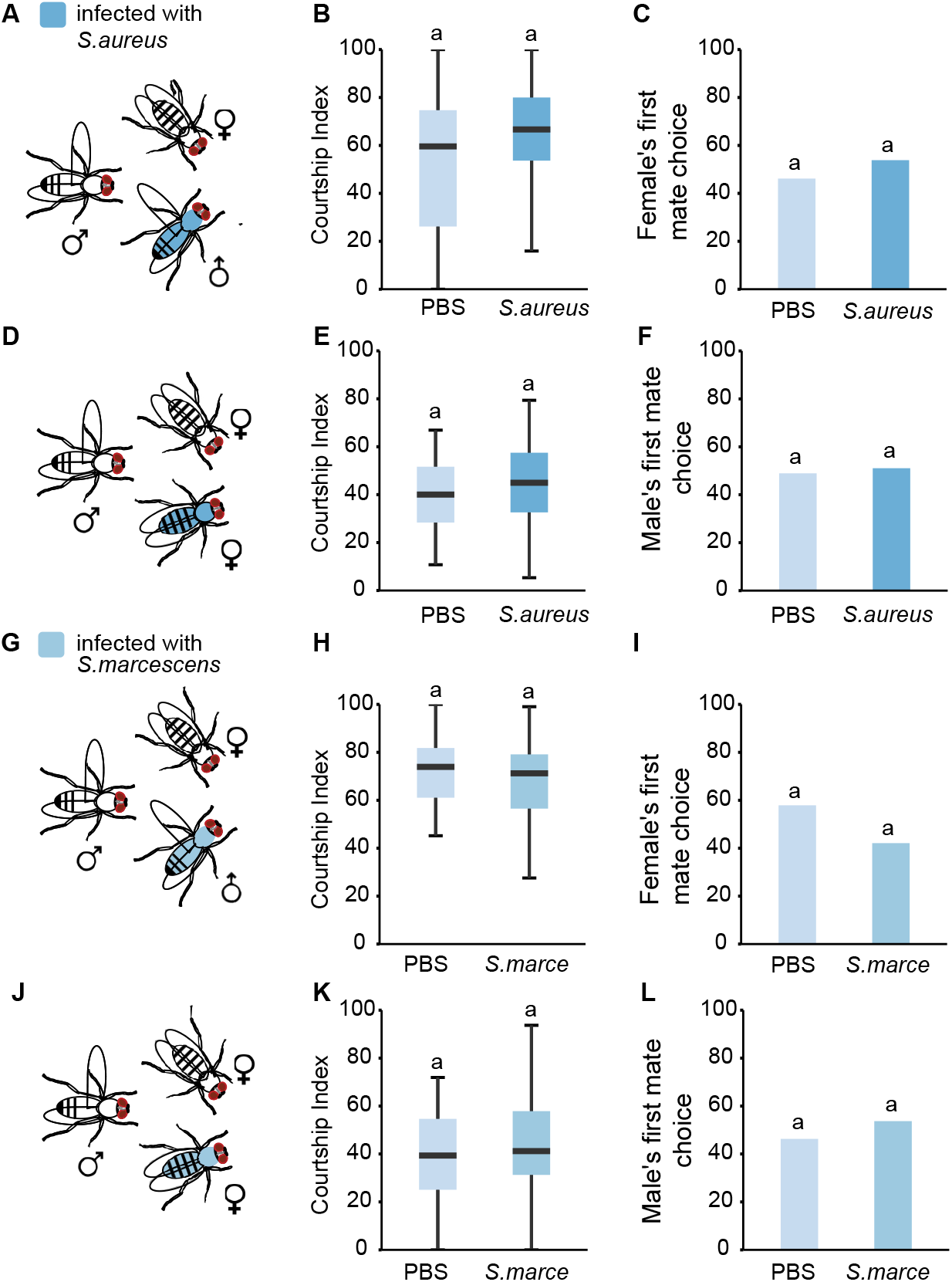
Effect of virulent infections on courtship behaviours and mate selection in a choice context. (A) A focal male was given a choice between a female injected with PBS and a female infected with *S. aureus*. (B) Courtship Index of the focal male towards either PBS or infected female (C) Male’s first mate choice (n=45). (D) A focal female was given a choice between a PBS male and a male infected with *S. aureus*. (E) Courtship Index of the PBS or infected male (F) Female’s first mate choice (n= 58). (G) A focal male was given a choice between a PBS female and a female infected with *S. marcescens*. (H) Courtship Index of the focal male towards either PBS or infected female (I) Male’s first mate choice (n=55). (J) A focal female was given a choice between a PBS male and a male infected with *S. marcescens*. (K) Courtship Index of the PBS or infected male (L) Female’s first mate choice (n=38). Wilcoxon test in B, E, H and K and Fisher Test in C, F, I and L. No significant differences were observed between the treatments.

### Activation of the immune system has limited impact on male courtship behaviour and female sexual receptivity

Our findings show that infections with several bacterial species that activate both arms of the immune system do not alter male or female reproductive behaviours in flies. We next wondered if induction of the immune system without the presence of bacteria would affect the reproductive behaviours shown by flies. To trigger a strong activation of each of the immune arms in a specific and controlled manner, we genetically induced the *Drosophila* immune pathways in the fat body, the site of humoral immune response, using the GAL4/UAS system. To achieve this, we overexpressed UAS-Imd [52] or UAS-Toll10B [53] using C564-GAL4 to target the fat body [54]. We found that activation of the Imd pathway did not significantly alter the courtship index or mating success in *C564>Imd* males (Median Courtship Index, *C564-GAL4/+* = 86, *UAS-Imd/+* = 93.8 and *C564-GAL4 > Uas-Imd* = 85.5, **Fig 6 A-B**), (Mating success, *C564-GAL4/+* = 90%, UAS IMD/+ = 89.47% and *C564-GAL4 > UAS-Imd* = 70%, **Fig 6C**). Similarly, activation of the Imd pathway did not influence female copulation latency (Median Copulation Latency, *C564-GAL4/+* =253.4s, *UAS-Imd/+* = 149.09s and *C564-GAL4>UAS-Imd* = 209.78, **Fig 6 G-H** or mating success (Mating success, *C564-GAL4/+* =100%, *UAS-Imd/+* = 85% and *C564-GAL4>UAS-Imd* = 100%, **Fig 6I**). Since the expression of Toll10B inhibits growth and causes developmental defects [56], we expressed Toll10B in an adult-specific way. We combined C564-GAL4 with the temperature-sensitive GAL80 (GAL80^TS^) [55], an inhibitor of GAL4. At high temperature, GAL80 ceases to suppress GAL4, thereby allowing the expression of Toll10B. Here again, we found that adult specific activation of Toll10B did not affect the courtship index or mating success in *C564-GAL4; GAL80^TS^>UAS-Toll10B* males when compared to the controls (Median Courtship Index, *C564-GAL4; tub-GAL80^TS^/+* = 95.68, *UAS-Toll10B/+* = 94.93 and *C564-GAL4; tub-GAL80^TS^>UAS-Toll10B* =98.56, **Fig 6D-E**), (Mating Success, *C564-GAL4; tub-GAL80^TS^/+*= 84.21%, *UAS-Toll10B/+* = 90.47% and *C564-GAL4; tub-GAL80^TS^>UAS-Toll10B* = 100%, **Fig 6F**). Furthermore, female sexual behaviour was unaffected in *C564-GAL4;GAL80^TS^>UAS-Toll10B females* (**Fig 6J-L**). These results, together with our previous findings, demonstrate that the activation of the immune system does not impact pre-copulatory behaviours in *Drosophila*.

**Figure 6.**
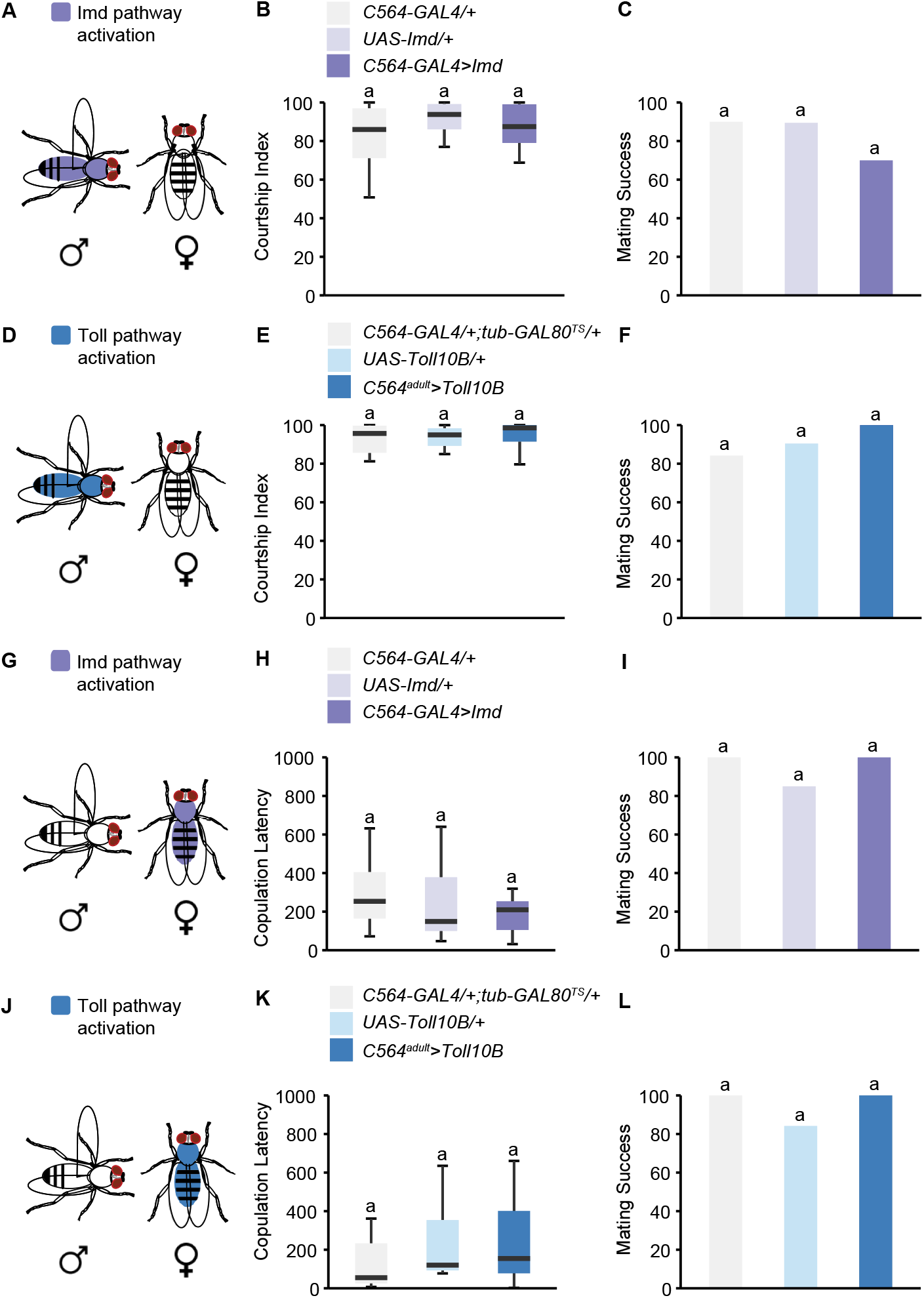
Effect of immune system activation on male courtship behaviour. (A, D) The Imd or the Toll pathway was artificially activated in the fat body of male flies. (B-C) Courtship index and mating success of *C564-GAL4>UAS-Imd* male flies (n=19-20). (E-F) Courtship index and mating success of *C564 GAL4; tub-GAL80^TS^>UAS-Toll10B* male flies (n=19-21). Effect of immune system activation on female sexual receptivity. (G, J) Imd or the Toll pathway were artificially activated in fat bodies of virgin female flies. (H, I) Copulation latency and mating success of *C564-GAL4>UAS-Imd* female flies (n=20-24). (K, L) Copulation latency and mating success of *C564-GAL4; tub-GAL80TS>UAS-Toll10B* female flies (n=19-20). Dunn Test in B, E, H and K, Fisher test IN C, F, I and L. No significant differences were observed between the genotypes.

## Discussion

Life history theory argues that there is a trade-off between energetically expensive traits, like reproduction and immunity [2,58,59]. Animals expend a considerable amount of energy in pre-copulatory traits, such as courtship behaviours to woo a potential mate [29]. At the same time, when challenged with an infection, individuals must allocate resources in mounting an effective immune response [56,57]. This involves fulfilling demands for protein synthetic pathways necessary for fighting infections and repairing damaged tissues. How do individuals prioritise and balance their investment in reproduction and immune defence? The trade-off between immunity and reproduction has been shown to be context-dependent and affected by a multitude of factors, including the genetic background of the host, its internal state, and the nature of the pathogen [20,53]. Yet, how these external and internal factors influence reproductive and defence strategies remain unclear.

Here, we study the impact of bacterial infection on reproductive behaviours in the fruit fly *Drosophila*. To ensure a comprehensive analysis, we employ six bacterial pathogens that vary in their virulence, phylogeny, and host-pathogen relationship. We evaluate and describe diverse aspects of reproductive behaviours in response to infection using two different wild-type *Drosophila melanogaster* strains. By studying infected males, we measure mating success and courtship intensity towards the healthy females. In addition, we study copulation latency and mating success of infected females exposed to healthy males. Remarkably, our findings indicate that, regardless of the type of bacterial species, and host strain, infection does not significantly affect male or female reproductive behaviours. Specifically, our findings show that infected male flies do not modify their courtship intensity or mating success, even when presented with unfavourable targets. These observations are consistent with a previous study by Keesey et al, which showed that infection with the lethal pathogen *Pseuduomonas entomophila* led to a small decrease in mating success in flies [38]. *Drosophila* exhibits sexual dimorphisms in immune system responses [51]. However, just as in males, sexual receptivity and mating success remained unaffected in infected females, though we did observe increased latency to copulation in sham-infected females. Furthermore, constitutive activation of the *Drosophila* immune system had no influence on male or female pre-copulatory behaviours. Bacterial infections may affect the probability of an individual of being selected as a mate. However, we saw no evidence for preference for healthy flies or flies infected with virulent pathogens in mate selection assays.

Our finding that *Drosophila* pre-copulatory behaviours are preserved during infections have parallels in other species. For instance, Greenspan et al. reported that frogs infected with a deadly pathogen have comparable calling properties to that of uninfected individuals [58]. In addition, while certain cricket species show a reduction in pre-copulatory behaviours upon infection [59–61], the immune activation by LPS injection in male crickets *Teleogryllus commodus* or *Gryllodes sigillatus* does not impact pre-copulatory traits [62,63]. It can be argued that, in order to maintain pre-copulatory traits intact through the infection, immune challenged male and female flies invest more of their available resources in reproduction, supporting the idea that sick flies make a terminal investment [64]. However, the terminal investment hypothesis classically predicts that infected animals will increase their reproductive investment compared to healthy animals. While we observed a preservation of the pre-copulatory behaviours, we did not find an increased expression of these traits.

The insect fat body is analogous to vertebrate adipose tissue and liver and is the primary site of energy storage [65]. At the same time, this organ is the source of the transcriptionally induced humoral immune response [66]. Activation of immune responses by exposure to pathogens limits resource utilisation towards other important biological processes. In line with this, several of the treatments employed in this study, e.g., *L. monocytogenes* or *E. coli* infections and genetic activation of the Toll pathway in the fat body, have been shown to interfere with the insulin signalling pathway and subsequently decrease nutrient storage or growth [45,67,68]. Moreover, infections with *L. monocytogenes* and *E. coli*, as well as numerous pathogens, cause systemic changes in metabolic signalling [32,45,68,69]. Yet, despite the marked metabolic switch triggered during systemic infections, male and female flies retain their pre-copulatory behaviours. These findings highlight the relevance of reproductive behaviours and raise the question as to what mechanisms are in place to preserve them. Are the neuronal clusters or tissues dedicated to courtship insensitive to the systemic transformations triggered by infection? Conversely, do courtship circuits actively sense infection-induced signals and respond by triggering mechanisms that maintain behavioural performance? Future studies should address the molecular and cellular machineries that control pre-copulatory behaviours upon bacteria detection. Some of the challenges include determining if the neurons involved in different male and female pre-copulatory behaviours express components of immune signalling cascade, and whether pathogen detection leads to physiological changes (e.g., modulation of neural activity via changes in the expression of neurotransmitters, neuromodulators and/or ion channels receptors).

Successful reproduction requires the completion of different events, spanning from mate attraction and courtship through to sperm transfer, egg fertilisation and offspring production. These pre and post-copulatory traits may differ in their response to infections. Indeed, studies have shown that bacterial infections in flies affect several post-copulatory processes. For instance, *Drosophila* mated females reduce the rate of egg-laying in response to *E. coli* infection or exposure to bacterial cell wall components, likely decreasing the infection risk of their progeny [37]. Moreover, infection with *P. rettgeri* affects expression of genes involved in oogenesis [70]. *Drosophila* males infected with *P. aeruginosa*, on the other hand, display a decrease in sperm viability [71]. A reduced investment in post-copulatory traits but not in the execution of courtship behaviours could be explained by differential energetic costs associated with these traits. Post-copulatory traits, like sperm production, ovulation and oviposition, might be energetically more expensive and therefore be under stronger selection pressure.

In addition to post-copulatory traits, local and systemic responses triggered by infection have been shown to affect several non-reproductive behaviours in *Drosophila*. For instance, fruit flies exposed to infection modify their behaviour to avoid dangerous food. Similar to the sauce béarnaise syndrome, upon ingesting food contaminated with bacteria (e.g., *ECC15* or *Pseudomonas entomophila*), flies reduce their activity and avoid harmed food via conditioned taste aversion mechanisms [72]. Moreover, upon contacting chemicals that normally activate the immune system, flies increase hygienic grooming [73]. Further, there is an interplay between immune activation and locomotion, with a subsequent impact on sleep, which depends on the pathogen type and the context and life history of the host. For instance, infection with *E. coli* leads to enhanced morning sleep, a process that is mediated by signals from peripheral immune systems (e.g., fat body) and the circadian clock [74,75]. Recently, it was reported that the Toll pathway in the fat body mediates a decrease of sleep in males infected with *M. luteus* [76]. In contrast, flies infected with *S. pneumonia* show normal levels of activity but with an altered sleep architecture and circadian rhythms [77].

In conclusion, despite the profound impact of bacterial infections on numerous metabolic, physiological and behavioural traits in *Drosophila*, pre-copulatory behaviours remain preserved, even in the face of deadly pathogens. Future experiments will investigate the mechanisms and the evolutionary ramifications of such strategy prioritisation.

## Materials and Methods

### *Drosophila* stocks

Wild-type fly lines used in the study include the *D. melanogaster* Canton-S and Dahomey strains. Transgenic lines include C564-GAL4 (BDSC 6982), UAS-Toll10B (BDSC 58987) and UAS-Imd (gift from Markus Knaden). Flies were raised on a standard cornmeal diet (recipe in **Table S2**) at 25 °C, 50-60% humidity with a 12 h light/dark cycle. Virgin male and female flies were collected <1 day after eclosion and were aged for 5-7 days in same-sex groups of 15-20 before experimentation.

**Table S2.**
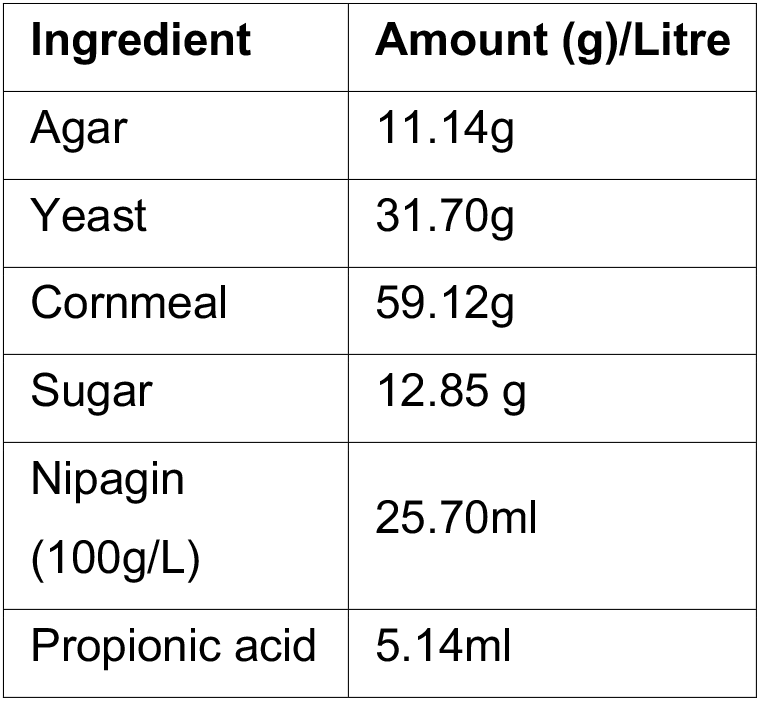
Fly food recipe.

| Ingredient | Amount (g)/Litre |
| --- | --- |
| Agar | 11.14g |
| Yeast | 31.70g |
| Cornmeal | 59.12g |
| Sugar | 12.85 g |
| Nipagin<br>(100g/L) | 25.70ml |
| Propionic acid | 5.14ml |

### Bacterial infection

The bacterial strains used in this study include *Serratia marcescens (DB11), Staphylococcus aureus (SH1000), Listeria monocytogenes* (EGD-e), *Escherichia coli (DH5α), Pectinobacterium carotovorum carotovorum 15 (ECC15)* and *Micrococcus luteus* (clinical isolate, gift from Prof William Wade, King’s College London). The bacterial strains were cultured overnight at conditions given below (**Table S3**). Cultures were pelleted down by centrifugation at 4500g for 2 minutes. The pellet was diluted in filter-sterilised PBS (Phosphate buffered saline) to a defined concentration (**Table S3**). 50 nl of diluted bacterial solution was injected employing a nano-injector (MPP1-3 Pressure Injector, Applied Scientific Instrumentation) into the abdomen of anaesthetised flies as previously described [42].

**Table S3.**
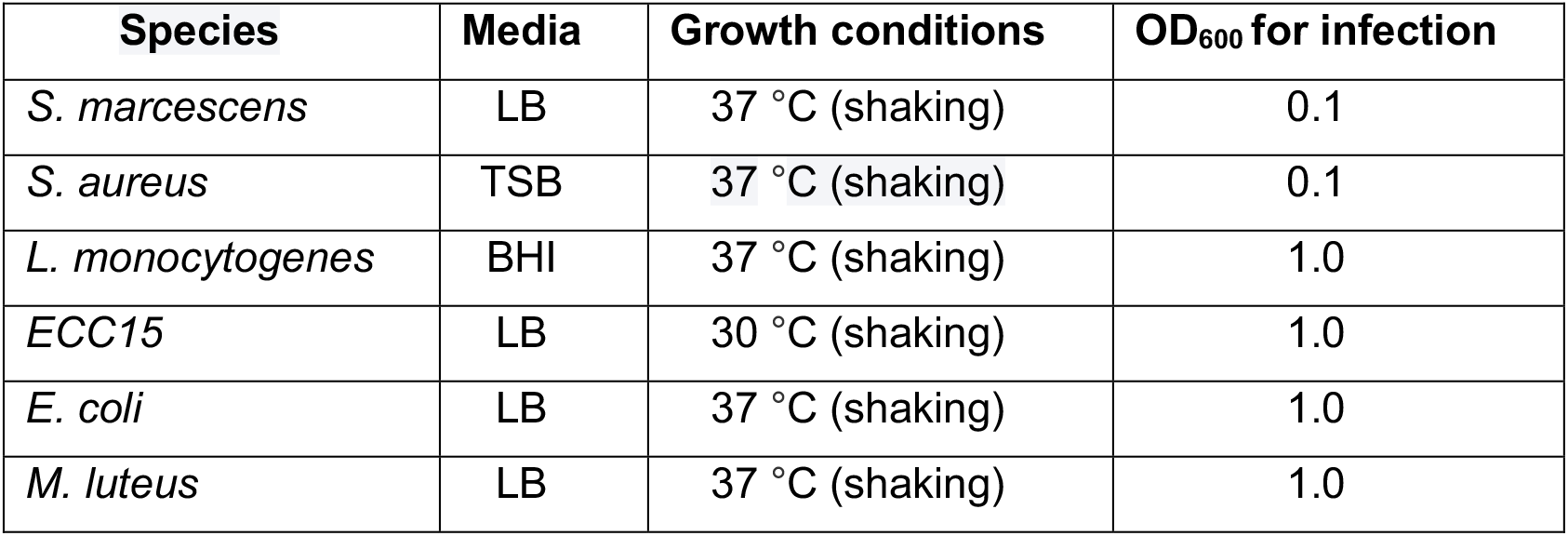
List of bacteria employed in the study.

| Species | Media | Growth conditions | OD<sub>600</sub> for infection |
| --- | --- | --- | --- |
| *S. marcescens* | LB | 37 °C (shaking) | 0.1 |
| *S. aureus* | TSB | 37 °C (shaking) | 0.1 |
| *L. monocytogenes* | BHI | 37 °C (shaking) | 1.0 |
| *ECC15* | LB | 30 °C (shaking) | 1.0 |
| *E. coli* | LB | 37 °C (shaking) | 1.0 |
| *M. luteus* | LB | 37 °C (shaking) | 1.0 |

### Survival assay

Infected flies and controls were placed in groups of 10-15 in vials at 29°C. The vials were inspected on an hourly/daily basis and the number of live flies were counted until all the flies were dead.

### Behavioural assays

All behavioural experiments were done in between zeitgeber time (ZT) 01 and ZT10 at 25°C. Mating assays were carried out in circular courtship chambers (20mm in diameter, 5 mm in height), which have built-in dividers that allow separation of the sexes before the experiment. The lighting was constant throughout the assay.

### Single pair mating assay

Flies were injected with bacteria or vehicle solution (PBS) and immediately placed them in the courtship chamber with food to avoid starvation. Before the behavioural measurement began, the uninfected flies of the opposite sex were introduced using a fly aspirator. The dividers were opened just before the assay and behaviours were recorded for one hour.

### Mate choice assays

In competitive mating assays, a focal fly was given a choice between an infected and a healthy (PBS) mate. The infected and healthy flies were marked with acrylic paint 48 hours before experimentation. After injection, both infected and PBS flies were transferred to courtship chamber with food. The focal fly was aspirated into the chamber before behavioural experimentation and behaviours were recorded for one hour.

For single pair mating assays, the following parameters were quantified: **courtship latency:** measured as the time taken by the male to initiate courtship, **courtship index:** measured as the proportion of time the male spends courting from the beginning of courtship until 10 minutes or end of copulation, **mating success:** measured as the number of successful copulations within one hour and **copulation latency:** measured as the time taken to copulate from the start of courtship. For competitive mating assays, the focal fly’s first mate choice was recorded: i.e if the fly chose to mate with a healthy or infected mate.

### Statistical analysis

All statistical analyses and data visualisation (ggplot2, R Markdown) were performed using R studio (R version 3.6). All data were tested for normality using Shapiro–Wilk test. As most of the behavioural data do not follow a normal distribution, Kruskal Wallis followed by Dunn test with Bonferroni corrections were used for post-hoc comparisons. Fisher’s exact test was used to analyse count data and log-rank test for survival data. Differences were accepted as significant at p <0.05.

## Data accessibility

This article has no additional data. R code used for analysis has been deposited at Github: https://github.com/salonirose95/Infection-paper/

## Authors’ contributions

Conceptualization, S.R., and C.R.; Methodology, S.R., E.J.B., R.C.M, M.D, and C.R.; Investigation, S.R., and C.B.; Resources, R.C.M, M.D, and C.R; Writing, S.R., E.J.B., R.C.M, M.D, and C.R.; Supervision, C.R.; Funding Acquisition, C.R.

## Competing interests

We declare we have no competing interests.

## Funding

This work was supported by grants from BBSRC (BB/S009299/1), Welcome Trust (214062/Z/18/Z) and Royal Society Research (RGS/R2/180272) to C.R, and a Darwin studentship to S.R. E. B. is supported by an IBRO Return Home Fellowship 2020.

## Acknowledgements

We thank Jean-Christophe Billeter, Joanne Yew, and Ana Depetris-Chauvin for helpful discussions. We are grateful to Bruno Lemaitre, Markus Knaden and Alicia Hidalgo for sharing fly stocks or bacterial strains with us. We would also like to thank Shaleen Glasgow for help with fly collections and Laurie Cazale-Debat for providing schematics. Finally, we thank the members of the Rezaval and Dionne lab for helpful discussions.

## Notes

### Competing Interest Statement

The authors have declared no competing interest.

## References

1. Schwenke RA, Lazzaro BP, Wolfner MF. 2016 Reproduction–Immunity Trade-Offs in Insects. Annu. Rev. Entomol. 61, 239–256. (doi:10.1146/annurev-ento-010715-023924)

2. Stearns SC. 1989 Trade-Offs in Life-History Evolution. Functional Ecology 3, 259. (doi:10.2307/2389364)

3. Armitage SAO, Thompson JJW, Rolff J, Siva-Jothy MT. 2003 Examining costs of induced and constitutive immune investment in Tenebrio molitor. J Evolution Biol 16, 1038–1044. (doi:10.1046/j.1420-9101.2003.00551.x)

4. Demas GE, Chefer V, Talan MI, Nelson RJ. 1997 Metabolic costs of mounting an antigen-stimulated immune response in adult and aged C57BL/6J mice. American Journal of Physiology-Regulatory, Integrative and Comparative Physiology 273, R1631–R1637. (doi:10.1152/ajpregu.1997.273.5.R1631)

5. Roberts JR, Souillard R, Bertin J. 2011 Avian diseases which affect egg production and quality. In Improving the Safety and Quality of Eggs and Egg Products, pp. 376–393. Elsevier. (doi:10.1533/9780857093912.3.376)

6. Ahmed AM, Hurd H. 2006 Immune stimulation and malaria infection impose reproductive costs in Anopheles gambiae via follicular apoptosis. Microbes and Infection 8, 308–315. (doi:10.1016/j.micinf.2005.06.026)

7. Ahmed AM, Baggott SL, Maingon R, Hurd H. 2002 The costs of mounting an immune response are reflected in the reproductive fitness of the mosquito *Anopheles gambiae*. Oikos 97, 371–377. (doi:10.1034/j.1600-0706.2002.970307.x)

8. Wigby S, Suarez SS, Lazzaro BP, Pizzari T, Wolfner MF. 2019 Sperm success and immunity. In Current Topics in Developmental Biology, pp. 287–313. Elsevier. (doi:10.1016/bs.ctdb.2019.04.002)

9. Chargé R, Saint Jalme M, Lacroix F, Cadet A, Sorci G. 2010 Male health status, signalled by courtship display, reveals ejaculate quality and hatching success in a lekking species. Journal of Animal Ecology (doi:10.1111/j.1365-2656.2010.01696.x)

10. Kolluru GR, Grether GF, Dunlop E, South SH. 2009 Food availability and parasite infection influence mating tactics in guppies (Poecilia reticulata). Behavioral Ecology 20, 131–137. (doi:10.1093/beheco/arn124)

11. Pélabon C, Borg ÅA, Bjelvenmark J, Barber I, Forsgren E, Amundsen T. 2005 Do microsporidian parasites affect courtship in two-spotted gobies? Marine Biology 148, 189–196. (doi:10.1007/s00227-005-0056-8)

12. Rushbrook BJ, Barber I. 2006 Nesting, courtship and kidney hypertrophy in Schistocephalus-infected male three-spined stickleback from an upland lake. J Fish Biology 69, 870–882. (doi:10.1111/j.1095-8649.2006.01164.x)

13. Gustafsson L, Nordling D, Andersson MS, Sheldon BC, Qvarnström A, Hamilton WD, Howard JC. 1994 Infectious diseases, reproductive effort and the cost of reproduction in birds. Philosophical Transactions of the Royal Society of London. Series B: Biological Sciences 346, 323–331. (doi:10.1098/rstb.1994.0149)

14. Adamo SA, Jensen M, Younger M. 2001 Changes in lifetime immunocompetence in male and female Gryllus texensis (formerly G. integer): trade-offs between immunity and reproduction. Animal Behaviour 62, 417–425. (doi:10.1006/anbe.2001.1786)

15. Ahtiainen JJ, Alatalo RV, Kortet R, Rantala MJ. 2005 A trade-off between sexual signalling and immune function in a natural population of the drumming wolf spider Hygrolycosa rubrofasciata. J Evolution Biol 18, 985–991. (doi:10.1111/j.1420-9101.2005.00907.x)

16. Stahlschmidt ZR, Rollinson N, Acker M, Adamo SA. 2013 Are all eggs created equal? Food availability and the fitness trade-off between reproduction and immunity. Funct Ecol 27, 800–806. (doi:10.1111/1365-2435.12071)

17. Velando A, Drummond H, Torres R. 2006 Senescent birds redouble reproductive effort when ill: confirmation of the terminal investment hypothesis. Proc. R. Soc. B 273, 1443–1448. (doi:10.1098/rspb.2006.3480)

18. An D, Waldman B. 2016 Enhanced call effort in Japanese tree frogs infected by amphibian chytrid fungus. Biol. Lett. 12, 20160018. (doi:10.1098/rsbl.2016.0018)

19. Clancy LM, Cooper AL, Griffith GW, Santer RD. 2017 Increased Male-Male Mounting Behaviour in Desert Locusts during Infection with an Entomopathogenic Fungus. Sci Rep 7, 5659. (doi:10.1038/s41598-017-05800-4)

20. Copeland EK, Fedorka KM. 2012 The influence of male age and simulated pathogenic infection on producing a dishonest sexual signal. Proc. R. Soc. B 279, 4740–4746. (doi:10.1098/rspb.2012.1914)

21. Farchmin PA, Eggert A-K, Duffield KR, Sakaluk SK. 2020 Dynamic terminal investment in male burying beetles. Animal Behaviour 163, 1–7. (doi:10.1016/j.anbehav.2020.02.015)

22. Duffield KR, Bowers EK, Sakaluk SK, Sadd BM. 2017 A dynamic threshold model for terminal investment. Behav Ecol Sociobiol 71, 185. (doi:10.1007/s00265-017-2416-z)

23. Hales KG, Korey CA, Larracuente AM, Roberts DM. 2015 Genetics on the Fly: A Primer on the *Drosophila* Model System. Genetics 201, 815–842. (doi:10.1534/genetics.115.183392)

24. Kazama H. 2015 Systems neuroscience in Drosophila: Conceptual and technical advantages. Neuroscience 296, 3–14. (doi:10.1016/j.neuroscience.2014.06.035)

25. Markow TA, Beall S, Castrezana S. 2012 The wild side of life: Drosophila reproduction in nature. Fly (Austin) 6, 98–101. (doi:10.4161/fly.19552)

26. Ellendersen BE, von Philipsborn AC. 2017 Neuronal modulation of D. melanogaster sexual behaviour. Current Opinion in Insect Science 24, 21–28. (doi:10.1016/j.cois.2017.08.005)

27. Spieth HT. 1966 Drosophilid mating behaviour: The behaviour of decapitated females. Animal Behaviour 14, 226–235. (doi:10.1016/S0003-3472(66)80076-3)

28. Agrawal S, Safarik S, Dickinson M. 2014 The relative roles of vision and chemosensation in mate recognition of Drosophila melanogaster. Journal of Experimental Biology 217, 2796–2805. (doi:10.1242/jeb.105817)

29. Mowles SL, Jepson NM. 2015 Physiological Costs of Repetitive Courtship Displays in Cockroaches Handicap Locomotor Performance. PLoS ONE 10, e0143664. (doi:10.1371/journal.pone.0143664)

30. Hoffmann JA. 2003 The immune response of Drosophila. Nature 426, 33–38. (doi:10.1038/nature02021)

31. Lemaitre B, Hoffmann J. 2007 The Host Defense of Drosophila melanogaster. Annual Review of Immunology 25, 697–743. (doi:10.1146/annurev.immunol.25.022106.141615)

32. Dionne M. 2014 Immune-metabolic interaction in *Drosophila*. Fly 8, 75–79. (doi:10.4161/fly.28113)

33. Dionne MS, Pham LN, Shirasu-Hiza M, Schneider DS. 2006 Akt and foxo Dysregulation Contribute to Infection-Induced Wasting in Drosophila. Current Biology 16, 1977–1985. (doi:10.1016/j.cub.2006.08.052)

34. Wagner C et al. 2021 Constitutive immune activity promotes JNK- and FoxO-dependent remodeling of Drosophila airways. Cell Reports 35, 108956. (doi:10.1016/j.celrep.2021.108956)

35. Clark RI et al. 2013 MEF2 Is an In Vivo Immune-Metabolic Switch. Cell 155, 435–447. (doi:10.1016/j.cell.2013.09.007)

36. Buchon N, Silverman N, Cherry S. 2014 Immunity in Drosophila melanogaster — from microbial recognition to whole-organism physiology. Nat Rev Immunol 14, 796–810. (doi:10.1038/nri3763)

37. Kurz CL, Charroux B, Chaduli D, Viallat-Lieutaud A, Royet J. 2017 Peptidoglycan sensing by octopaminergic neurons modulates Drosophila oviposition. eLife 6, e21937. (doi:10.7554/eLife.21937)

38. Keesey IW, Koerte S, Khallaf MA, Retzke T, Guillou A, Grosse-Wilde E, Buchon N, Knaden M, Hansson BS. 2017 Pathogenic bacteria enhance dispersal through alteration of Drosophila social communication. Nat Commun 8, 265. (doi:10.1038/s41467-017-00334-9)

39. Mansfield BE, Dionne MS, Schneider DS, Freitag NE. 2003 Exploration of host-pathogen interactions using Listeria monocytogenes and Drosophila melanogaster. Cell Microbiol 5, 901–911. (doi:10.1046/j.1462-5822.2003.00329.x)

40. Duneau D, Ferdy J-B, Revah J, Kondolf H, Ortiz GA, Lazzaro BP, Buchon N. 2017 Stochastic variation in the initial phase of bacterial infection predicts the probability of survival in D. melanogaster. eLife 6, e28298. (doi:10.7554/eLife.28298)

41. Nehme NT, Liégeois S, Kele B, Giammarinaro P, Pradel E, Hoffmann JA, Ewbank JJ, Ferrandon D. 2007 A model of bacterial intestinal infections in Drosophila melanogaster. PLoS Pathogens 3, e173. (doi:10.1371/journal.ppat.0030173)

42. Dionne MS, Ghori N, Schneider DS. 2003 *Drosophila melanogaster* Is a Genetically Tractable Model Host for *Mycobacterium marinum*. Infect Immun 71, 3540–3550. (doi:10.1128/IAI.71.6.3540-3550.2003)

43. Patrnogic J, Castillo JC, Shokal U, Yadav S, Kenney E, Heryanto C, Ozakman Y, Eleftherianos I. 2018 Pre-exposure to non-pathogenic bacteria does not protect Drosophila against the entomopathogenic bacterium Photorhabdus. PLoS ONE 13, e0205256. (doi:10.1371/journal.pone.0205256)

44. Irving P, Troxler L, Heuer TS, Belvin M, Kopczynski C, Reichhart J-M, Hoffmann JA, Hetru C. 2001 A genome-wide analysis of immune responses in Drosophila. Proceedings of the National Academy of Sciences 98, 15119–15124. (doi:10.1073/pnas.261573998)

45. Vincent CM, Dionne MS. 2021 Disparate regulation of IMD signaling drives sex differences in infection pathology in *Drosophila melanogaster*. Proc Natl Acad Sci USA 118, e2026554118. (doi:10.1073/pnas.2026554118)

46. Eleftherianos I, More K, Spivack S, Paulin E, Khojandi A, Shukla S. 2014 Nitric Oxide Levels Regulate the Immune Response of Drosophila melanogaster Reference Laboratory Strains to Bacterial Infections. Infect. Immun. 82, 4169–4181. (doi:10.1128/IAI.02318-14)

47. Bass TM, Grandison RC, Wong R, Martinez P, Partridge L, Piper MDW. 2007 Optimization of Dietary Restriction Protocols in Drosophila. The Journals of Gerontology: Series A 62, 1071–1081. (doi:10.1093/gerona/62.10.1071)

48. Needham AJ, Kibart M, Crossley H, Ingham PW, Foster SJ. 2004 Drosophila melanogaster as a model host for Staphylococcus aureus infection. Microbiology (Reading, Engl.) 150, 2347–2355. (doi:10.1099/mic.0.27116-0)

49. Ejima A, Smith BPC, Lucas C, van der Goes van Naters W, Miller CJ, Carlson JR, Levine JD, Griffith LC. 2007 Generalization of Courtship Learning in Drosophila Is Mediated by cis-Vaccenyl Acetate. Current Biology 17, 599–605. (doi:10.1016/j.cub.2007.01.053)

50. Belmonte RL, Corbally M-K, Duneau DF, Regan JC. 2020 Sexual Dimorphisms in Innate Immunity and Responses to Infection in Drosophila melanogaster. Front. Immunol. 10, 3075. (doi:10.3389/fimmu.2019.03075)

51. Nunn CL, Lindenfors P, Pursall ER, Rolff J. 2009 On sexual dimorphism in immune function. Phil. Trans. R. Soc. B 364, 61–69. (doi:10.1098/rstb.2008.0148)

52. Buchon N, Broderick NA, Poidevin M, Pradervand S, Lemaitre B. 2009 Drosophila Intestinal Response to Bacterial Infection: Activation of Host Defense and Stem Cell Proliferation. Cell Host & Microbe 5, 200–211. (doi:10.1016/j.chom.2009.01.003)

53. Maxton-Küchenmeister J, Handel K, Schmidt-Ott U, Roth S, Jäckle H. 1999 Toll homolog expression in the beetle Tribolium suggests a different mode of dorsoventral patterning than in Drosophila embryos. Mechanisms of Development 83, 107–114. (doi:10.1016/S0925-4773(99)00041-6)

54. Harrison DA, Binari R, Nahreini TS, Gilman M, Perrimon N. 1995 Activation of a Drosophila Janus kinase (JAK) causes hematopoietic neoplasia and developmental defects. The EMBO Journal 14, 2857–2865. (doi:10.1002/j.1460-2075.1995.tb07285.x)

55. del Valle Rodríguez A, Didiano D, Desplan C. 2012 Power tools for gene expression and clonal analysis in Drosophila. Nat Methods 9, 47–55. (doi:10.1038/nmeth.1800)

56. Cotter SC, Simpson SJ, Raubenheimer D, Wilson K. 2011 Macronutrient balance mediates trade-offs between immune function and life history traits. Functional Ecology 25, 186–198. (doi:10.1111/j.1365-2435.2010.01766.x)

57. Lochmiller RL, Deerenberg C. 2000 Trade-offs in evolutionary immunology: just what is the cost of immunity? Oikos 88, 87–98. (doi:10.1034/j.1600-0706.2000.880110.x)

58. Greenspan SE, Roznik EA, Schwarzkopf L, Alford RA, Pike DA. 2016 Robust calling performance in frogs infected by a deadly fungal pathogen. Ecol Evol 6, 5964–5972. (doi:10.1002/ece3.2256)

59. Jacot A, Scheuber H, Brinkhof MWG. 2004 COSTS OF AN INDUCED IMMUNE RESPONSE ON SEXUAL DISPLAY AND LONGEVITY IN FIELD CRICKETS. Evolution 58, 2280–2286. (doi:10.1111/j.0014-3820.2004.tb01603.x)

60. Jacot A, Scheuber H, Kurtz J, Brinkhof MWG. 2005 Juvenile immune status affects the expression of a sexually selected trait in field crickets. J Evolution Biol 18, 1060–1068. (doi:10.1111/j.1420-9101.2005.00899.x)

61. Tregenza T, Simmons LW, Wedell N, Zuk M. 2006 Female preference for male courtship song and its role as a signal of immune function and condition. Animal Behaviour 72, 809–818. (doi:10.1016/j.anbehav.2006.01.019)

62. Drayton JM, Boeke JEK, Jennions MD. 2013 Immune Challenge and Pre- and Post-copulatory Female Choice in the Cricket Teleogryllus commodus. J Insect Behav 26, 176–190. (doi:10.1007/s10905-012-9347-3)

63. Kerr AM, Gershman SN, Sakaluk SK. 2010 Experimentally induced spermatophore production and immune responses reveal a trade-off in crickets. Behavioral Ecology 21, 647–654. (doi:10.1093/beheco/arq035)

64. Clutton-Brock TH. 1984 Reproductive Effort and Terminal Investment in Iteroparous Animals. The American Naturalist 123, 212–229. (doi:10.1086/284198)

65. Arrese EL, Soulages JL. 2010 Insect Fat Body: Energy, Metabolism, and Regulation. Annu. Rev. Entomol. 55, 207–225. (doi:10.1146/annurev-ento-112408-085356)

66. Hoffmann JA. 2003 The immune response of Drosophila. Nature 426, 33–38. (doi:10.1038/nature02021)

67. DiAngelo JR, Bland ML, Bambina S, Cherry S, Birnbaum MJ. 2009 The immune response attenuates growth and nutrient storage in Drosophila by reducing insulin signaling. Proceedings of the National Academy of Sciences 106, 20853–20858. (doi:10.1073/pnas.0906749106)

68. Chambers MC, Song KH, Schneider DS. 2012 Listeria monocytogenes Infection Causes Metabolic Shifts in Drosophila melanogaster. PLoS ONE 7, e50679. (doi:10.1371/journal.pone.0050679)

69. Vincent CM, Simoes da Silva CJ, Wadhawan A, Dionne MS. 2020 Origins of Metabolic Pathology in Francisella-Infected Drosophila. Front. Immunol. 11, 1419. (doi:10.3389/fimmu.2020.01419)

70. Short SM, Lazzaro BP. 2013 Reproductive Status Alters Transcriptomic Response to Infection in Female *Drosophila melanogaster*. G3 Genes|Genomes|Genetics 3, 827–840. (doi:10.1534/g3.112.005306)

71. Radhakrishnan P, Fedorka KM. 2012 Immune activation decreases sperm viability in both sexes and influences female sperm storage. Proc. R. Soc. B. 279, 3577–3583. (doi:10.1098/rspb.2012.0654)

72. Kobler JM, Rodriguez Jimenez FJ, Petcu I, Grunwald Kadow IC. 2020 Immune Receptor Signaling and the Mushroom Body Mediate Post-ingestion Pathogen Avoidance. Current Biology 30, 4693–4709.e3. (doi:10.1016/j.cub.2020.09.022)

73. Yanagawa A, Guigue AMA, Marion-Poll F. 2014 Hygienic grooming is induced by contact chemicals in Drosophila melanogaster. Front. Behav. Neurosci. 8. (doi:10.3389/fnbeh.2014.00254)

74. Williams JA, Sathyanarayanan S, Hendricks JC, Sehgal A. 2007 Interaction Between Sleep and the Immune Response in Drosophila: A Role for the NFκB Relish. Sleep 30, 389–400. (doi:10.1093/sleep/30.4.389)

75. Kuo T-H, Pike DH, Beizaeipour Z, Williams JA. 2010 Sleep triggered by an immune response in Drosophila is regulated by the circadian clock and requires the NFkappaB Relish. BMC Neurosci 11, 17. (doi:10.1186/1471-2202-11-17)

76. Vincent CM, Beckwith EJ, Pearson WH, Kierdorf K, Gilestro G, Dionne MS. 2021 Infection increases activity via *Toll* dependent and independent mechanisms in *Drosophila melanogaster*. (doi:10.1101/2021.08.24.457493)

77. Shirasu-Hiza MM, Dionne MS, Pham LN, Ayres JS, Schneider DS. 2007 Interactions between circadian rhythm and immunity in Drosophila melanogaster. Current Biology 17, R353–R355. (doi:10.1016/j.cub.2007.03.049)

