## Supplementary information for "Pre-copulatory reproductive behaviours are preserved in *Drosophila melanogaster* infected with bacteria"

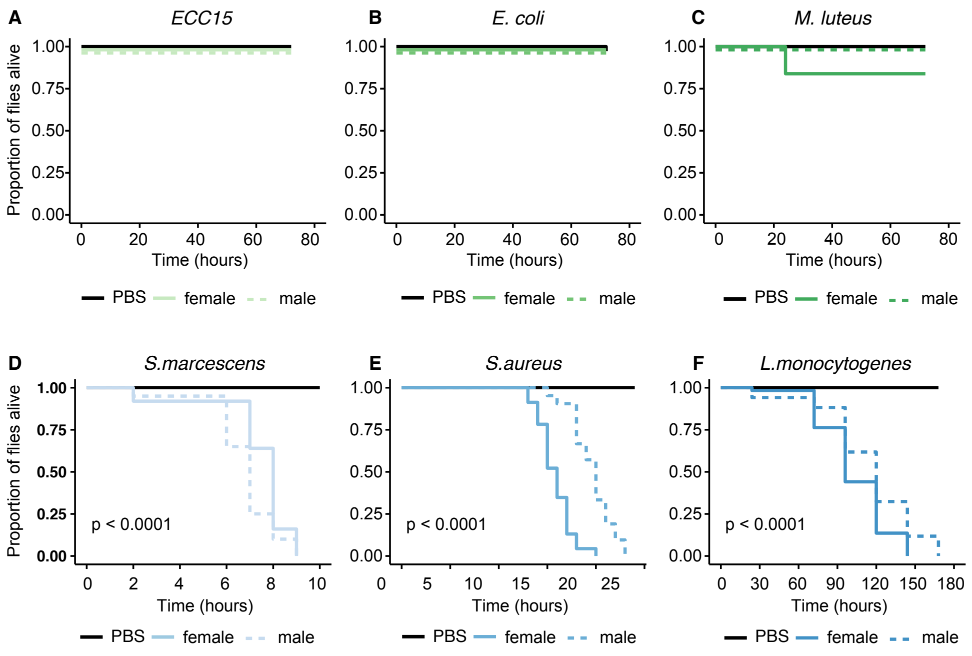

**Supplementary Figure 1**. Effect of septic infection on survival on wild-type flies. CS flies were injected with 50 nl of bacterial solution or PBS and the number of survivors was quantified at regular intervals. (A) *ECC15* (n = 15-40), (B) *E. coli* (n = 20-42) and (C) *M. luteus* (n=19-41) and infection did not reduce host survival, p>0.05. (D) *S. marcescens* (n= 20), (E) *S. aureus* (n = 20-21) and (F) *L. monocytogenes* (n=20-34) are lethal pathogens and kill 100% of the hosts. Statistical analysis was performed with the log-rank test.

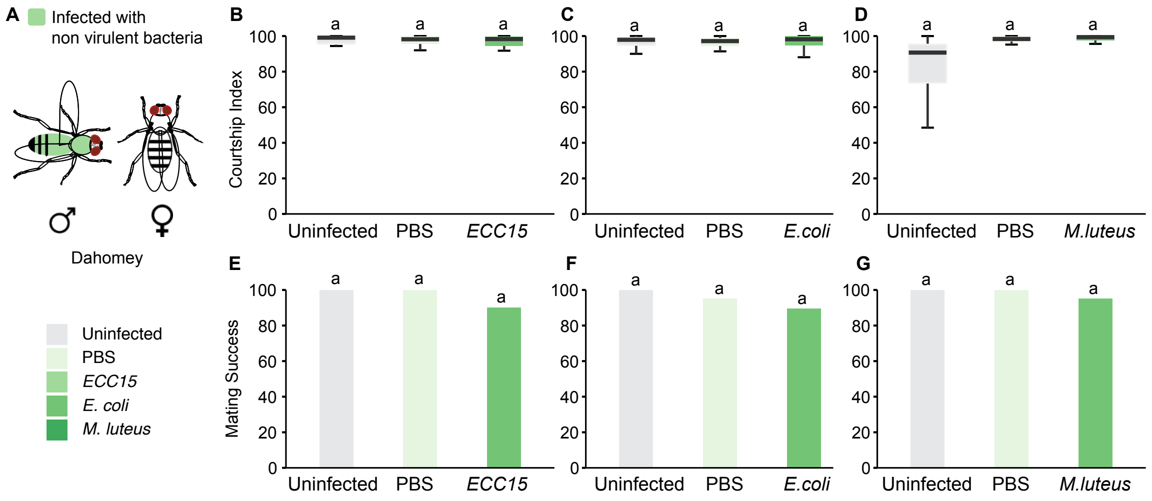

**Supplementary Figure 2.** Effect of non-pathogenic bacterial infections on male courtship behaviour. (A) Male Dahomey flies were injected with three different pathogens and tested in a single pair courtship assay with an uninfected virgin female. (B) Courtship index and (C) mating success of males infected with *ECC15* and their respective controls (n=19-21). (D) Courtship index and (E) mating success of males infected with *E. coli* and their respective controls (n=18-20). (F) Courtship index and (G) mating success of males infected with *M. luteus* and their respective controls (n=20-22). Dunn Test in B-D and Fisher Test in E-G. No significant differences were observed between the treatments.

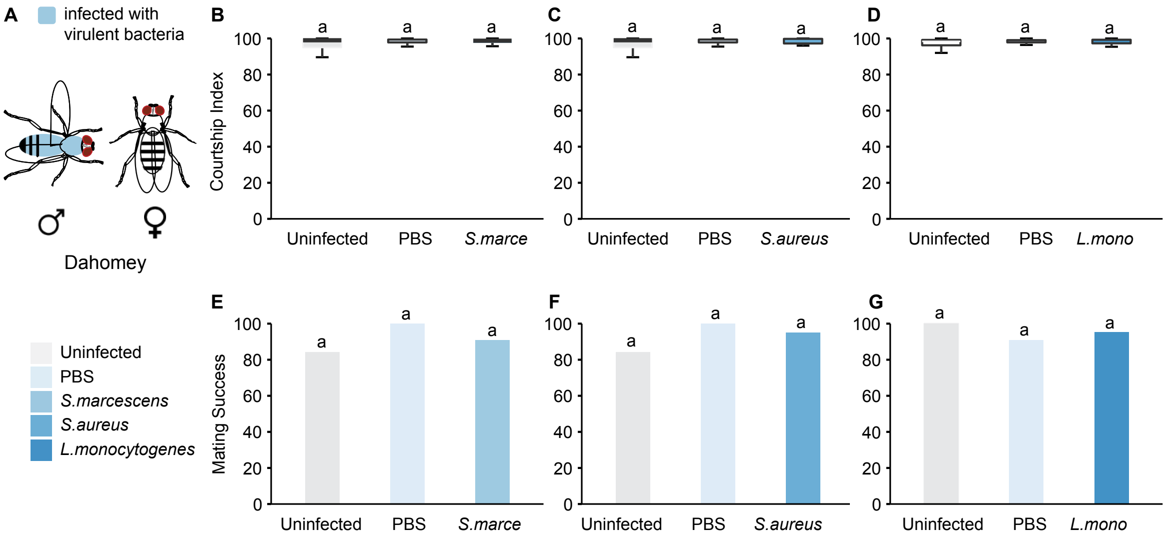

**Supplementary Figure 3.** Effect of pathogenic bacterial infections on male courtship behaviour. (A) Male Dahomey flies were injected with three different pathogens and tested in a single pair courtship assay with an uninfected virgin female. (B) Courtship index and (C) mating success of males infected with *S. marcescens* and their respective controls (n=20-22). (D) Courtship index and (E) mating success of males infected with *S. aureus* and their respective controls (n=19-21). (F) Courtship index and (G) mating success of males infected with *L. monocytogenes* and their respective controls (n=21-22). Dunn Test in B-D and Fisher Test in E-G. No significant differences were observed between the treatments.

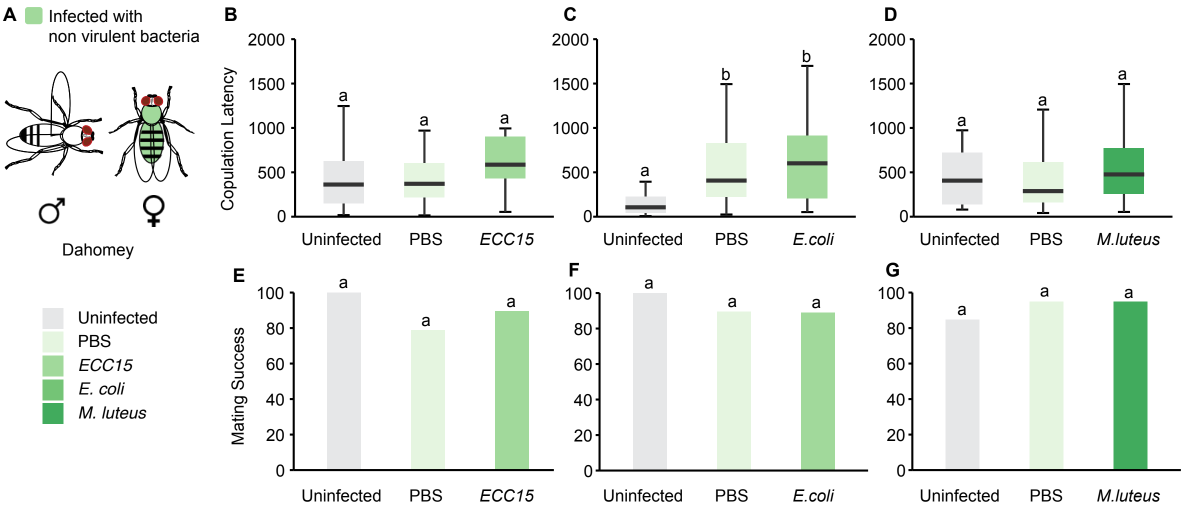

**Supplementary Figure 4.** Effect of non-virulent bacterial infections on female receptivity. Virgin female Dahomey flies were injected with three different pathogens and tested in a single pair courtship assay with an uninfected male. (A) Copulation latency and (D) mating success of females infected with *ECC15* and their respective controls (n=18-20). (B) Copulation latency and (E) mating success of females infected with *E. coli* and their respective controls (n=19-21). (C) Copulation latency and (F) mating success of males infected with *M. luteus* and their respective controls (n=13-20). Dunn Test in B-D and Fisher Test in E-G.

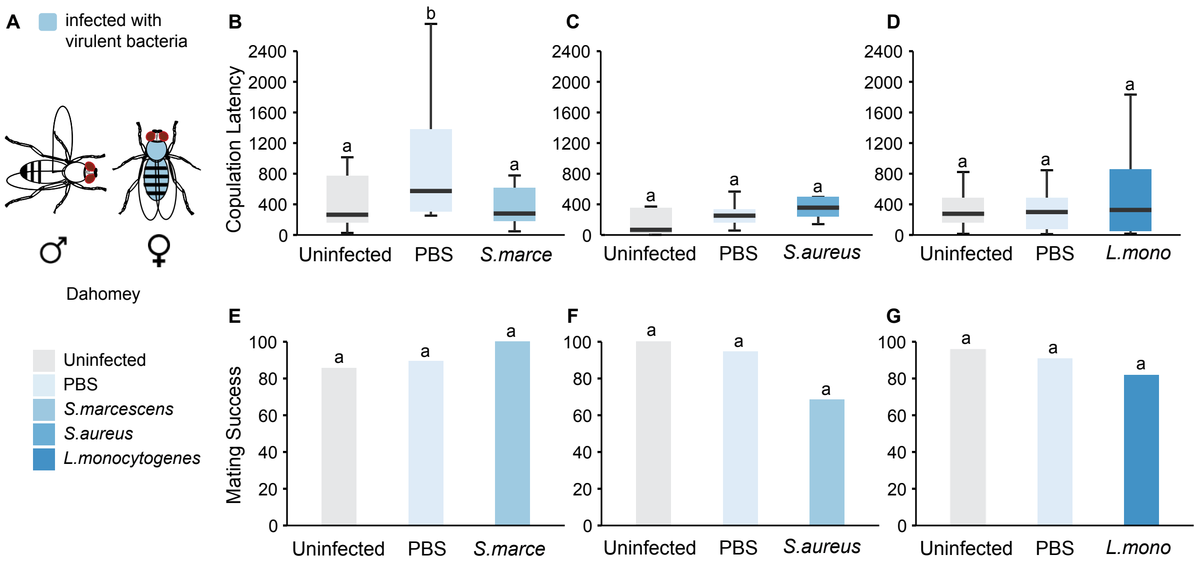

**Supplementary Figure 5**: Effect of virulent bacterial infections on female receptivity. (A) Virgin Dahomey female flies were injected with three different pathogens and tested in a single pair courtship assay with an uninfected male. (B) Copulation latency and (E) mating success of females infected with *S. marcescens* and their respective controls (n=19). (C) Copulation latency and (F) mating success of females infected with *S. aureus* and their respective controls (n=22-23). (D) Copulation latency and (G) mating success of males infected with *L. monocytogenes* and their respective controls (n=19-20). Dunn Test in B-D and Fisher Test in E-G.

| Experiment 1 | | | |
|---|---|---|---|
| **Infection status** | **Median courtship index** | **IQR** | **Sample size** |
| Uninfected | 10 | 14.65 | 36 |
| PBS | 14.7 | 16.2 | 27 |
| *M. luteus* | 18.75 | 31.325 | 36 |
| **Comparison** | **P value** | | |
| *M. luteus* - PBS | 0.9676892 | | |
| *M. luteus* - Uninfected | 0.1424981 | | |
| PBS - Uninfected | 0.855051 | | |
| **Experiment 2** | | | |
| **Infection status** | **Median courtship index** | **IQR** | **Sample size** |
| Uninfected | 20.9 | 23.9 | 20 |
| PBS | 28.6 | 40.5 | 33 |
| *S. marcescens* | 12.4 | 32.3 | 34 |
| **Comparison** | **P value** | | |
| *S. marcescens* - PBS | 0.2264451 | | |
| *S. marcescens* - Uninfected | 1 | | |
| PBS - Uninfected | 1 | | |
| **Experiment 3** | | | |
| **Infection status** | **Median courtship index** | **IQR** | **Sample size** |
| Uninfected | 9.72 | 8.3 | 16 |
| PBS | 17.5 | 18.4 | 14 |
| *ECC15* | 34.3 | 35.5 | 22 |
| **Comparison** | **P value** | | |
| *ECC15* - PBS | 0.69586157 | | |
| *ECC15* - Uninfected | 0.00192666 | | |
| PBS - Uninfected | 0.01493624 | | |

**Table S1**. Effect on bacterial infections on male courtship behaviour. Male CS flies were injected with three different pathogens and tested in a single pair courtship assay with an uninfected mated female. Their median courtship index and inter-quartile range is reported along with post-hoc pairwise comparisons using Dunn Test.
